# Suppression of liquid-liquid phase separation by 1,6-hexanediol partially compromises the 3D genome organization in living cells

**DOI:** 10.1101/2020.05.18.101261

**Authors:** Sergey V. Ulianov, Artem K. Velichko, Mikhail D. Magnitov, Artem V. Luzhin, Arkadiy K. Golov, Natalia Ovsyannikova, Igor I. Kireev, Alexander V. Tyakht, Alexey A. Gavrilov, Omar L. Kantidze, Sergey V. Razin

## Abstract

Liquid-liquid phase separation (LLPS) contributes to the spatial and functional segregation of molecular processes. However, the role played by LLPS in chromatin folding in living cells remains unclear. Here, using stochastic optical reconstruction microscopy (STORM) and Hi-C techniques, we studied the effects of 1,6-hexanediol (1,6-HD)-mediated LLPS modulation on higher-order chromatin organization in living cells. We found that 1,6-HD treatment caused the enlargement of nucleosome nanodomains and their more uniform distribution in the nuclear space. At a megabase-scale, chromatin underwent moderate but irreversible perturbations that resulted in the partial mixing of A and B compartments. The removal of 1,6-HD from the culture medium did not allow chromatin to acquire initial configurations, but increased further mixing of the chromatin compartments and resulted in more compact repressed chromatin than in untreated cells. 1,6-HD treatment also weakened enhancer-promoter interactions but did not considerably affect CTCF-dependent loops. Our results suggest that 1,6-HD-sensitive LLPS plays a limited role in chromatin spatial organization by constraining its folding patterns and facilitating compartmentalization at different levels.

## INTRODUCTION

The modern concept of hierarchical chromatin folding in the eukaryotic cell nucleus is based on the results of Hi-C analyses (Lieberman-Aiden et al. 2009). Eukaryotic chromosomes are partitioned into semi-independent topologically associating domains (TADs) (Dixon et al. 2012; Nora et al. 2012), which are composed of chromatin loops (Rao et al. 2014). Low-resolution analyses have demonstrated that active and repressed chromatin are spatially segregated into A and B chromatin compartments, respectively (Lieberman-Aiden et al. 2009), which are comprised of smaller compartmental domains (Rowley et al. 2017). Recent evidence suggests that the basic spatial organization of the genome relies on the interplay between active DNA loop extrusion and the passive spatial segregation of chromatin domains enriched in particular sets of epigenetic marks (such as active and inactive chromatin domains) (Rowley et al. 2017; Schwarzer et al. 2017; Nuebler et al. 2018; Rowley and Corces 2018). Although the DNA loop extrusion machinery has been properly characterized (Fudenberg et al. 2017; Davidson et al. 2019), the processes/forces that mediate the spatial segregation of active and inactive chromatin remain poorly understood. Recent studies have disclosed the important role played by liquid-liquid phase separation (LLPS) in the functional compartmentalization of the eukaryotic cell nucleus, particularly in the assembly of various nuclear bodies, such as the nucleolus, splicing speckles, and Cajal bodies (Courchaine et al. 2016; Banani et al. 2017; Boeynaems et al. 2018; Gomes and Shorter 2019). The proteins that participate in the formation of phase-separated condensates frequently possess intrinsically disordered regions (IDRs) (Meng et al. 2015; Uversky 2017a; Darling et al. 2018). These IDRs may mediate weak-affinity and non-specific interactions with multiple target sites that trigger the LLPS (Uversky 2017b). IDRs are present in many proteins involved in the assembly of repressive chromatin domains (histone H1 (Turner et al. 2018), heterochromatin protein one (HP1) (Larson et al. 2017) and chromobox 2 (CBX2) subunit of mammalian PRC1 complex (Tatavosian et al. 2019)), which enables these factors to form liquid condensates, both *in vitro* and *in vivo* (Strom et al. 2017; Turner et al. 2018; Tatavosian et al. 2019). On the other hand, the components of the transcriptional machinery, including RNA polymerase II (Boehning et al. 2018), Mediator complex subunits (Nagulapalli et al. 2016), and various transcription factors (Boija et al. 2018), also possess IDRs and are capable of forming complex phase-separated condensates at enhancers (Cho et al. 2018; Sabari et al. 2018). The assembly of activating domains at enhancers and the clustering of RNA polymerase II molecules at transcription hubs or factories via phase separation appears to be functionally relevant (Hnisz et al. 2017; Hahn 2018; Sabari et al. 2018; Gurumurthy et al. 2019; Nair et al. 2019). Therefore, the formation of different types of phase-separated chromatin condensates may underlie the segregation of the A and B chromatin compartments (Rada-Iglesias et al. 2018). In addition to the direct LLPS-driven segregation of chromatin domains bearing different epigenetic marks, the spatial clustering of active and repressed chromatin domains may also be mediated by their differential interactions with nuclear bodies, such as nucleoli and nuclear speckles (Quinodoz et al. 2018). The disruption of nuclear speckles was shown to reduce spatial chromatin interactions within the active compartment (Hu et al. 2019). Considering that nucleoli and nuclear speckles are LLPS-dependent membrane-free compartments (Uversky 2017a), LLPS is likely to contribute to the spatial segregation of A and B chromatin compartments, both directly and indirectly.

To obtain further insights into the possible roles played by LLPS in 3-dimensional (3D) genome organization, here, we studied the effects of 1,6-hexanediol (1,6-HD), an agent known to disrupt liquid-phase condensates (Sabari et al. 2018), on chromatin folding in HeLa cells. Using stochastic optical reconstruction microscopy (STORM) and Hi-C analysis, we demonstrate that the suppression of LLPS leads to the partial decondensation of compact nucleosome nanodomains, irreversibly changes the internal structures of A and B chromatin compartments, and slightly compromises their spatial segregation. At the level of TADs, 1,6-HD-driven alteration of LLPS changes chromatin compaction and weakens enhancer-promoter loops. Upon the restoration of LLPS after 1,6-HD removal the repressed chromatin did not return to its initial state but instead acquires a novel, more compact configuration.

## RESULTS

### 1,6-HD treatment of living cells to study higher-order chromatin organization

To investigate the role played by LLPS in spatial genome organization, we used human HeLa cells that were treated with 1,6-HD. This aliphatic alcohol is predicted to disrupt weak hydrophobic interactions, both *in vitro* and *in vivo*, disassembling LLPS-dependent macromolecular condensates (Kroschwald et al. 2017; Alberti et al. 2019; Lesne et al. 2019). Molecular condensates depended purely on phase separation driven by electrostatic interactions are expected to be unaffected by 1,6-HD treatment (Lesne et al. 2019). Nevertheless, 1,6-HD is the only available tool to test *in vivo* the contribution of LLPS in the assembly of macromolecular assemblies at this moment (Kroschwald et al. 2017). 1,6-HD has been used in a number of seminal studies to treat living cells, particularly the studies that described the involvement of LLPS in super-enhancer function and heterochromatin domain formation (Larson et al. 2017; Cho et al. 2018; Sabari et al. 2018). Prolonged treatment with 1,6-HD can lead to cellular membrane rupture and cell death (Kroschwald et al. 2017; Ming et al. 2019). To minimize 1,6-HD toxicity *in vivo* and to avoid possible secondary effects associated with such toxicity, we utilized the following precautions: i) limited 5% 1,6-HD treatment to 15 minutes; and ii) applied 1,6-HD to HeLa cells that were first transiently permeabilized with Tween 20 (Michelini et al. 2019). Treatment of living cells with 5% 1,6-HD is widely accepted condition (Molliex et al. 2015; Yamazaki et al. 2018; Oshidari et al. 2020; Zhang et al. 2020) that provides disruption of LLPS-driven condensates with minimal cytotoxicity. Using cells transiently permeabilized with Tween 20 additionally lowers 1,6-HD toxicity that in case of intact non-permeabilized cells is related to the cytoplasmic membrane rupture (Kroschwald et al. 2017; Ming et al. 2019). Generally, the short-term treatment with Tween 20 to permeabilize eukaryotic cells does not affect cell morphology or metabolic processes, such as transcription and DNA repair foci formation (Francia et al. 2012; Michelini et al. 2019). Additionally, we have verified that the treatments used in our study did not result in cell death and did not perturb intracellular processes. The treatment of HeLa cells with Tween 20 alone (1%, 10 min) did not lead to the appearance of apoptotic cells and did not inhibit RNA polymerase I and II-dependent transcription (Fig. 1A and B). The short-term incubation of transiently permeabilized (Tween 20-treated) cells in culture medium containing 1,6-HD (5%, 15 min) induced only moderate levels of apoptosis (Fig. 1A). Moreover, the number of apoptotic cells did not significantly increase during the 1.5-h recovery period, during which cells were incubated in Tween 20- and 1,6-HD-free medium (Fig. 1A). Although transcription was strongly inhibited by 1,6-HD treatment, it was almost fully restored after the recovery period (Fig. 1B).

**Figure. 1.**
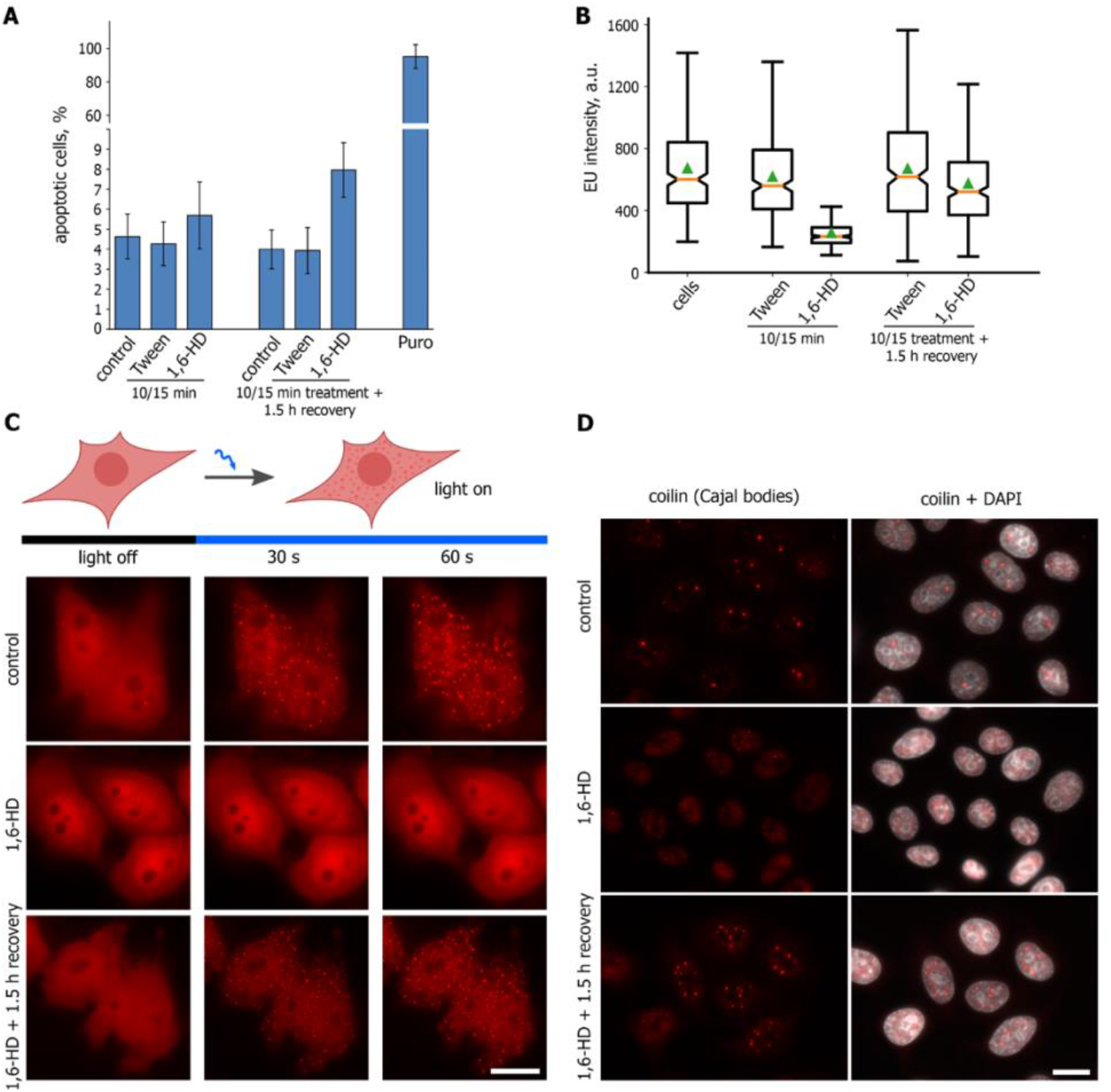
1,6-HD compromises LLPS in living human cells. (**A**) HeLa cells were untreated or treated with Tween 20 (1%, 10 min; “Tween”), Tween 20 followed with 1,6-HD (5%, 15 min; “1,6-HD”) before being analyzed using CellEvent Caspase 3/7 Detection Reagent. The cells treated as described and then incubated in fresh culture medium for 1.5 h were analysed as well. Percent of caspase 3/7-positive (apoptotic) cells is shown. (**B**) HeLa cells treated as described in (**A**) were pulsed with 5-ethyniluridine (EU, 200 μM, 15 min). Box plots show the EU fluorescence intensities. Horizontal lines represent the medians. (**C**) HeLa cells transfected with pHR-FUSN-mCh-Cry2WT were transiently permeabilized and then either mock-treated (control), treated with 1,6-HD (5%, 15 min), or treated with 1,6-HD and allowed to recover for 1.5 h. OptoDroplet formation was monitored as described in (Shin et al. 2017). (**D**) Transiently permeabilized HeLa cells were untreated (control), treated with 1,6-HD (5%, 15 min), or treated with 1,6-HD and allowed to recover for 1.5 h before being stained for coilin (red).

To test the *in vivo* LLPS-disrupting properties of 1,6-HD treatment, we applied two different approaches. First, we investigated the influence of 1,6-HD on LLPS using a recently developed optoDroplet system (Shin et al. 2017). In this system, the intrinsically disordered region of an RNP granule protein FUS is combined with the fluorescent protein mCherry and the light-sensitive oligomerization domain of *Arabidopsis thaliana* cryptochrome 2 (CRY2) to generate a fusion protein that undergoes LLPS in living cells upon blue light activation (Shin et al. 2017). HeLa cells, transfected with a plasmid encoding the above-described light-sensitive chimeric protein, were subjected to blue light irradiation in culture medium either with or without 1,6-HD (5%) (Fig. 1C). In contrast to Tween 20, 1,6-HD treatment strongly affected the assembly of optoDroplets in HeLa cells (Fig. 1C). After the 1.5-h-long incubation of the treated cells in 1,6-HD-free medium, the ability of the transfected cells to assemble optoDroplets in response to blue light irradiation was completely reestablished (Fig. 1C). Second, we examined whether the LLPS-dependent membrane-free intranuclear compartments, such as Cajal bodies and splicing speckles, were affected by 1,6-HD treatment. These compartments were choosen because they were reported to be sensitive to 1,6-HD (Lin et al. 2016; Guo et al. 2019). The indirect immunofluorescence analysis of coilin, a major proteinaceous component of Cajal bodies, showed that 1,6-HD treatment (5%, 15 min), but not the treatment with Tween 20 alone, disrupted these compartments in living cells (Fig. 1D and Supplemental Fig. S1A). During the 1.5-h-long recovery period in 1,6-HD-free medium, Cajal bodies were fully re-established (Fig. 1D and Supplemental Fig. S1A). Structured illumination microscopy (SIM) of SC35 (SRSF2) combined with the nanodomain clustering analysis (Ricci et al. 2015) also demonstrated that the treatment with 1,6-HD affected the integrity of splicing speckles (Supplemental Fig. S1B and C). These data examining 1,6-HD toxicity and its mode of action demonstrate the applicability of 1,6-HD treatments to the study of the role of LLPS in higher-order chromatin organization in eukaryotic cells.

### Chromatin domains became enlarged and more uniformly distributed upon 1,6-HD treatment

First, we analyzed the distribution of histone H2B in HeLa cells treated with or without 1,6-HD, using conventional epifluorescence microscopy (Supplemental Fig. S1C). Virtually no changes were detected in the distributions of H2B in 1,6-HD-treated cells compared with control cells. To examine the organization of chromatin at a nanoscale resolution, we performed STORM imaging of the core histone H2B in HeLa cells treated with or without 1,6-HD. In agreement with the previously published data (Ricci et al. 2015; Nozaki et al. 2017; Xu et al. 2018), H2B appeared to be clustered into discrete chromatin domains (Fig. 2A), which reflected the organization of nucleosomes into chromatin fibers (Ricci et al. 2015). Visual inspection of images suggested that 1,6-HD treatment made the chromatin domains distribution pattern more coarse. To analyze quantitatively the effects of 1,6-HD on chromatin organization in interphase nuclei, we applied the following two types of analyses: i) nanodomain clustering analyses (Ricci et al. 2015), and ii) spatial descriptive statistics (radial distribution function (RDF) and L-function (Nozaki et al. 2017; Xu et al. 2018)). Nanodomain clustering analyses were performed, as described previously (Ricci et al. 2015) (Fig. 2B). Quantitative analysis demonstrated a slight decrease in the number of histone H2B nanodomains identified in 1,6-HD-treated HeLa cells compared with control cells (Fig. 2C). Simultaneously, H2B nanodomain areas shifted to higher values in 1,6-HD-treated cells (Fig. 2D and E). Almost no change in the distributions of H2B nanodomain nearest neighbor distances was observed between control HeLa cells and cells treated with 1,6-HD (Fig. 2F).

**Figure. 2.**
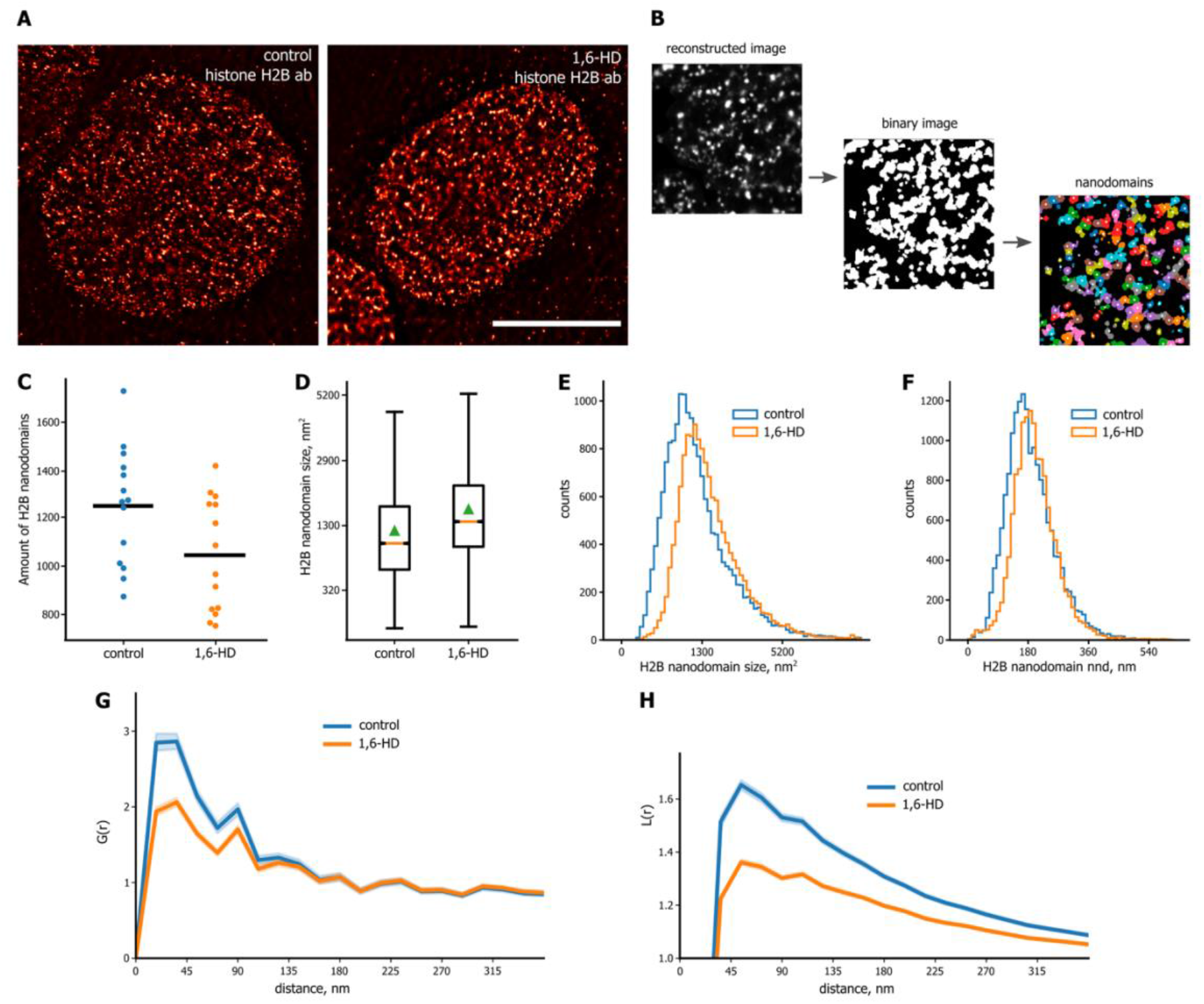
Super-resolution microscopy analysis of chromatin organization changes induced by 1,6-HD. (**A**) Representative STORM images of H2B in human HeLa cells that were either untreated (control) or treated with 1,6-HD (5%, 15 min). Scale bar: 5 μm. (**B**) Schematic representation of the analysis performed to cluster H2B nanodomains (details are provided in the Methods section). (**C-D**) Swarm and box plots demonstrating the numbers of H2B nanodomains (**C**) and their sizes (**D**) in HeLa cells treated as described in (**A**). On the swarm plots, horizontal lines represent the average values; on the box plots, horizontal lines represent the medians, triangles show the average values, the upper and lower ends of the boxplots show the upper and lower quartiles, and the whiskers indicate the upper and lower fences. (**E-F**) Distributions of H2B nanodomain sizes (**E**) and H2B nanodomain nearest neighbor distances (**F**) for cells treated as described in (**A**). (**G-H**) R- [G(r); **G**] and L-function (**H**) plots of chromatin using the same conditions as described in (**A**). For each condition, n = 10-15 cells.

RDF, G(r), shows the probability of identifying a molecule with respect to its neighboring molecules as a function of radial distance (r). The width of the RDF graph reflects the correlation length, and the height of the graph indicates the relative degree of clustering. RDF remains constant at all radial distances for randomly distributed molecules (Nozaki et al. 2017; Xu et al. 2018). In control HeLa cells, the RDF for histone H2B shows a narrow sharp peak suggesting the presence of highly clustered small chromatin domains (Fig. 2G), which is in a good agreement with the published data (Nozaki et al. 2017). 1,6-HD treatment resulted in reduced levels of H2B clustering (Fig. 2G). These data, when plotted as an L-function (L(r) versus r), clearly showed that the 1,6-HD treatment of HeLa cells resulted in H2B-marked chromatin domains becoming more uniformly distributed than in control cells (Fig. 2H).

Together, the analyses performed demonstrated that the disruption of LLPS in living cells led to the enlargement and more uniform distribution of chromatin domains in the nuclear space.

### Treatment of living cells with 1,6-HD affects chromatin compartment strength

To obtain further insights into the role played by LLPS in 3D genome organization, we performed *in situ* Hi-C analysis on HeLa cells that were first transiently permeabilized with Tween 20 and then either i) not treated (Control), ii) treated with 5% 1,6-HD for 15 min (Hex5), or iii) treated with 1,6-HD for 15 min and then allowed to recover in a fresh culture medium that did not contain neither Tween 20 nor 1,6-HD for 1.5 h (Recovery). Hi-C analysis was performed in two biological replicates, using the DpnII restriction enzyme. Each Hi-C library was sequenced to approximately 150 million paired-end reads per replicate, and 75-98 million unique contacts were obtained after data processing (Supplemental Table S1). Because the biological replicates were highly correlated (Supplemental Fig. S3A), we combined them to obtain 158-193 million sequenced ligation junctions per experiment (Supplemental Table S1), which allowed us to construct Hi-C maps at up to 20 kb resolution (Fig. 3A, Supplemental Fig. S2). Interestingly, the dependence of contact probability on genomic distance, *P_c_*(*s*), was virtually the same in the Control and Hex5 samples, but differed in the Recovery sample (Fig. 3B and Supplemental Fig. S4A), which was characterized by decreased spatial interactions over distances shorter than 107 bp and increased spatial interactions at longer distances. The ratio of *cis* (intrachromosomal) to *trans* (interchromosomal) contacts decreased approximately 2.5 times in the Recovery sample, although the ratios were similar between the Control and Hex5 samples (Fig. 3C, Supplemental Fig. S4B).

**Figure. 3.**
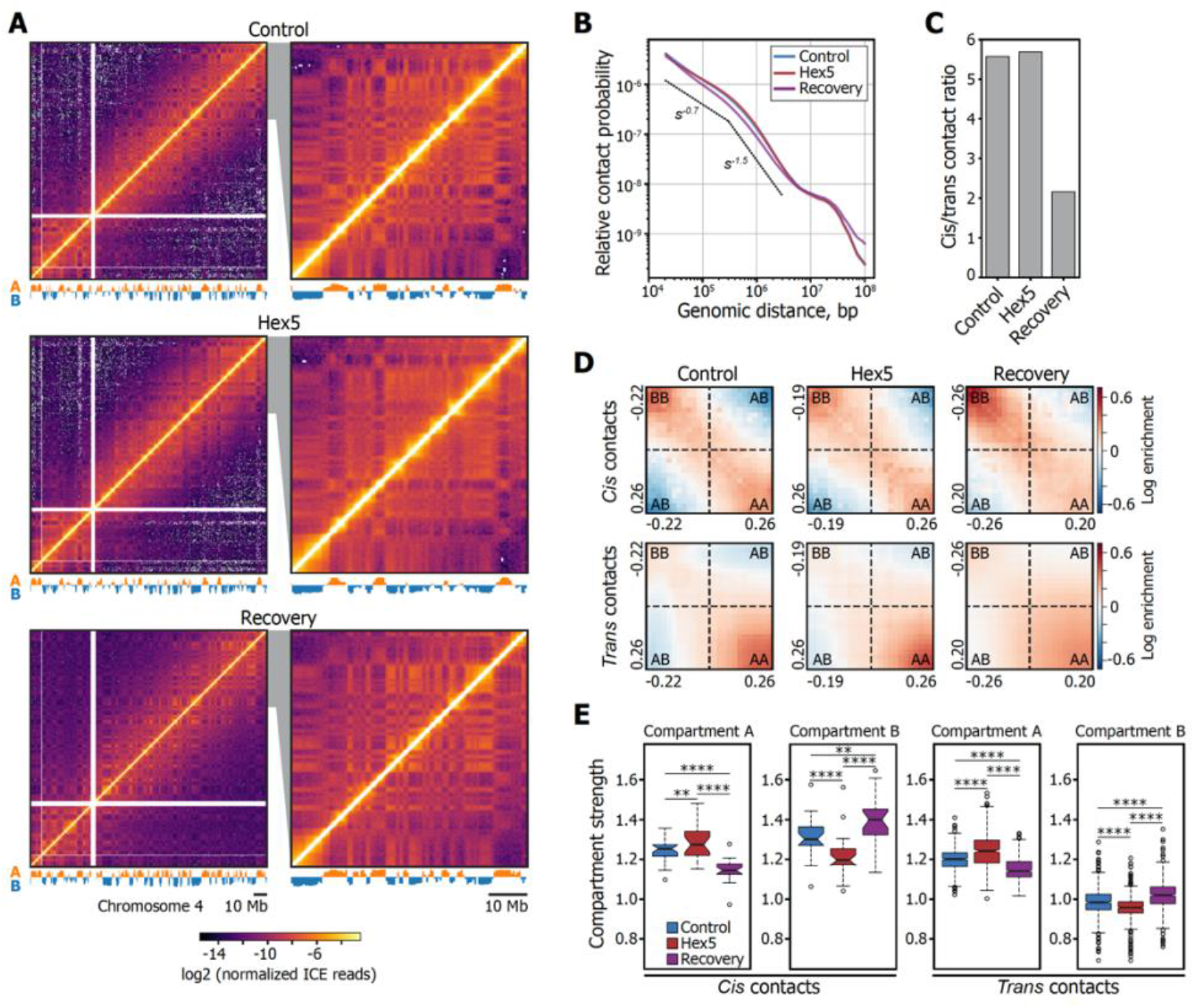
1,6-HD treatment alters the strengths of A and B compartments, both *in cis* and *in trans*. **(A)** Visualization of Control, Hex5 and Recovery contact matrices at a 500-kb resolution. The compartment profile is shown below the maps. (**B**) Dependence of contact probability, *P_c_*(*s*), on genomic distance, *s*, for Control, Hex5, and Recovery samples. Black lines show the slopes for *P_c_*(*s*) = *s-0.75* and *P_c_*(*s*) = *s-1.5*. (**C**) Ratio between *cis* (intrachromosomal) and *trans* (interchromosomal) contacts. (**D**) Heatmaps (saddle plots) showing log2 values of contact enrichments between genomic regions belonging to A (positive PC1 values) and B (negative PC1 values) compartments. Maximal and minimal values of PC1 are shown. (**E**) Compartment strength *in cis* and *in trans*. \*\*\*\**P* < 0.0001, \*\**P* < 0.01 in a Mann-Whitney U-test.

Visual inspection of the Hi-C heat maps revealed that the plaid pattern became less pronounced for the Hex5 sample compared with the Control or Recovery heat maps (Fig. 3A) indicating that the disruption of LLPS compromised chromatin compartmentalization. To systematically analyze the effects of 1,6-HD treatment on the A/B compartments, we performed independent principal component analyses (PCA) (Lieberman-Aiden et al. 2009) for the Control, Hex5, and Recovery contact matrices and used publicly available profiles of epigenetic markers (Supplemental Table S2) to validate compartment segmentation (Supplemental Fig. S3C). Because the values of the principal component 1 (PC1) were highly correlated between the Control, Hex5, and Recovery samples (Pearson’s correlation, *R* > 0.9; Supplemental Fig. 3D and F) and only approximately 8% of the genomic bins switched their compartment states between the Hex5 and Control samples (Supplemental Fig. S3E and F), we concluded that 1,6-HD treatment did not substantially change the A/B-compartment profile along the genome. We next separately analyzed the compartment strength (Falk et al. 2019) in the Control, Hex5, and Recovery matrices for *cis*- and *trans*-interactions. Interestingly, for all samples, interactions in *trans* occurred preferentially within the A compartment, whereas interactions in *cis* were more prominent in the B compartment (Fig. 3D and E). For both in *cis* and in *trans* the strength of the A compartment increased upon treatment with 1,6-HD and decreased to below the initial level in the Recovery sample. In contrast, the strength of the B compartment decreased in the Hex5 sample and increased to above the Control level in the Recovery sample. Thus, we observed opposing trends for the responses of the A and B compartments to LLPS disruption and subsequent recovery, which may indicate that different molecular mechanisms are responsible for forming each compartment.

To verify that transient permeabilization of the cells with Tween 20 did not contribute to the changes in higher-order chromatin organization observed, we performed additional *in situ* Hi-C analysis on HeLa cells either i) untreated, ii) treated with 1% Tween 20 for 10 min, or iii) treated with Tween 20 for 10 min and then allowed to recover 1h 45 min in a fresh culture medium that did not contain Tween 20. Hi-C analysis was performed in two biological replicates; sequencing statistics are shown in Supplemental Table S3. Analyses of Pearson’s correlation coefficient between the samples and the ratio of *cis* (intrachromosomal) and *trans* (interchromosomal) contacts clearly showed that the differences between these three samples were negligible (Supplemental Fig. S5A and B). The same conclusion was drawn based on the results of chromatin compartment mapping and analysis (Supplemental Fig. S5C-E). These results showed that the treatment of cells with Tween 20 did not alter the spatial genome organization. Therefore, the changes of the 3D genome observed in HeLa cells transiently permeabilized with Tween 20 and treated with 1,6-HD, can be attributed solely to the effects of 1,6-HD-mediated LLPS modulation.

### 1,6-HD treatment irreversibly modifies the internal structure of compartments and compromises their spatial segregation

To gain a more comprehensive picture of the intra- and intercompartment interactions that occur at various genomic distances, we developed a pentad analysis (see Online Methods). In this approach, we divided the entire Hi-C map for each chromosome into five characteristic fields, which formed a pentad that was averaged genome-wide (Fig. 4A). The A and B types lie on the map diagonal and represent interactions within continuous genomic fragments from the A and B compartments. Analyses of these zones allowed us to track contact frequencies at short distances inside compartments. Zones AA, BB, and AB do not lean on the map diagonal and represent long-range interactions both inside the compartments (AA and BB) and between them (AB). The pentads clearly showed that the distributions of spatial contacts within zones A and B were strikingly different (Fig. 4B). For all samples, the observed contact frequency at the map diagonal was higher than expected for the A compartment and was lower than expected for the B compartment, which was enriched with more distant interactions. This result reflects the more compact chromatin state in the B compartment.

**Figure. 4.**
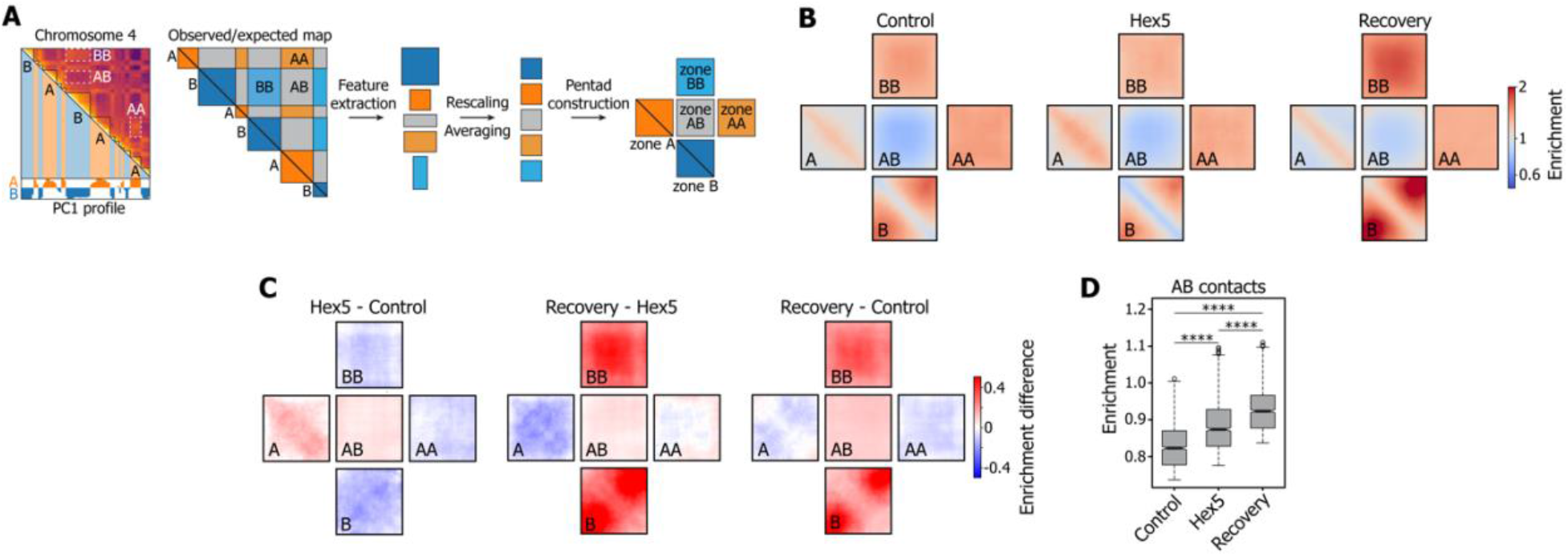
1,6-HD treatment compromises the spatial segregation between the A and B compartments. (**A**) Schematic representation of the pentad analysis. (**B**) Plots (pentads) showing the observed over expected contact frequencies inside the A and B compartments at short (A, B) and large genomic distances (AA, BB) and between compartments (AB). (**C**) Pairwise subtractions of Control, Hex5, and Recovery pentads. (**D**) Boxplots showing the contact frequencies between the A and B compartments. \*\*\*\**P* < 0.0001 in a Mann-Whitney U-test.

Comparisons between the Control and Hex5 pentads showed that the above-described total increase in the A compartment strength observed for the Hex5 sample (Fig. 3D and E) was entirely mediated by the gain of contacts at short distances (zone A), whereas long-range interactions (zone AA) were depleted in 1,6-HD-treated cells (Fig. 4C). In contrast, the overall decrease in the B compartment strength observed in the Hex5 sample was mediated by the depletion of spatial contacts at both short- and long-range distances. These changes in the intracompartmental contact profiles following 1,6-HD treatment were accompanied by increased interactions between compartments, indicating the weakening of their spatial segregation (Fig. 4C and D). Strikingly, removing 1,6-HD from the cell media (Recovery) did not restore the initial contact frequencies, either inside compartments or between them. In the Recovery sample, compared with the Control sample, the B compartment became much more compact, both at short- and long-range distances, whereas the number of contacts inside the entire A compartment generally decreased. We also observed the further mixing of the A and B compartments following 1,6-HD removal. Taken together, these results suggested that 1,6-HD treatment removed some constrains on large-scale chromatin folding, which resulted in perturbations that could not be easily reversed but allowed for the alternative refolding of chromatin fiber during the recovery experiment. Previously, Amat et al. (Amat et al. 2019) reported hyperosmotic shock-induced changes in chromatin compartments that closely resembled the changes observed in our study, as demonstrated by the analysis of their data using the pentad algorithm (Supplemental Fig. S6). However, hyperosmotic shock-induced changes were fully reversed upon the return of cells to normal conditions.

### 1,6-HD treatment reversibly changes TAD compaction and weakens enhancer-promoter loops

We next analyzed the effects of 1,6-HD treatment on chromatin folding, at the scale of TADs and loops. In the Hex5 sample, compared with the Control sample, TADs became more compact in the A compartment and less compact in the B compartment (Fig. 5A). This result is in agreement with the observed changes in contact frequencies at short genomic distances inside both compartments (Fig. 4C). In contrast to the contact frequencies in the compartments, the contact frequencies inside TADs almost fully reverted to the Control level in the Recovery sample. Because mammalian TADs are formed by CTCF/cohesin-mediated loop extrusion (Sanborn et al. 2015; Fudenberg et al. 2016; Mirny et al. 2019), we assumed that 1,6-HD treatment did not affect the CTCF binding profile across the entire genome, which allowed chromatin fibers to adopt their initial compaction profiles inside TADs after 1,6-HD removal. To test this hypothesis, we performed chromatin immunoprecipitation with anti-CTCF antibodies, followed by DNA sequencing (ChIP-seq). Indeed, the CTCF binding profiles in the Control and Hex5 samples were virtually identical (Fig. 5B and C, Supplemental Fig. S7). Accordingly, the strength of CTCF-mediated loops only slightly decreased upon 1,6-HD treatment (Fig. 5D).

**Figure. 5.**
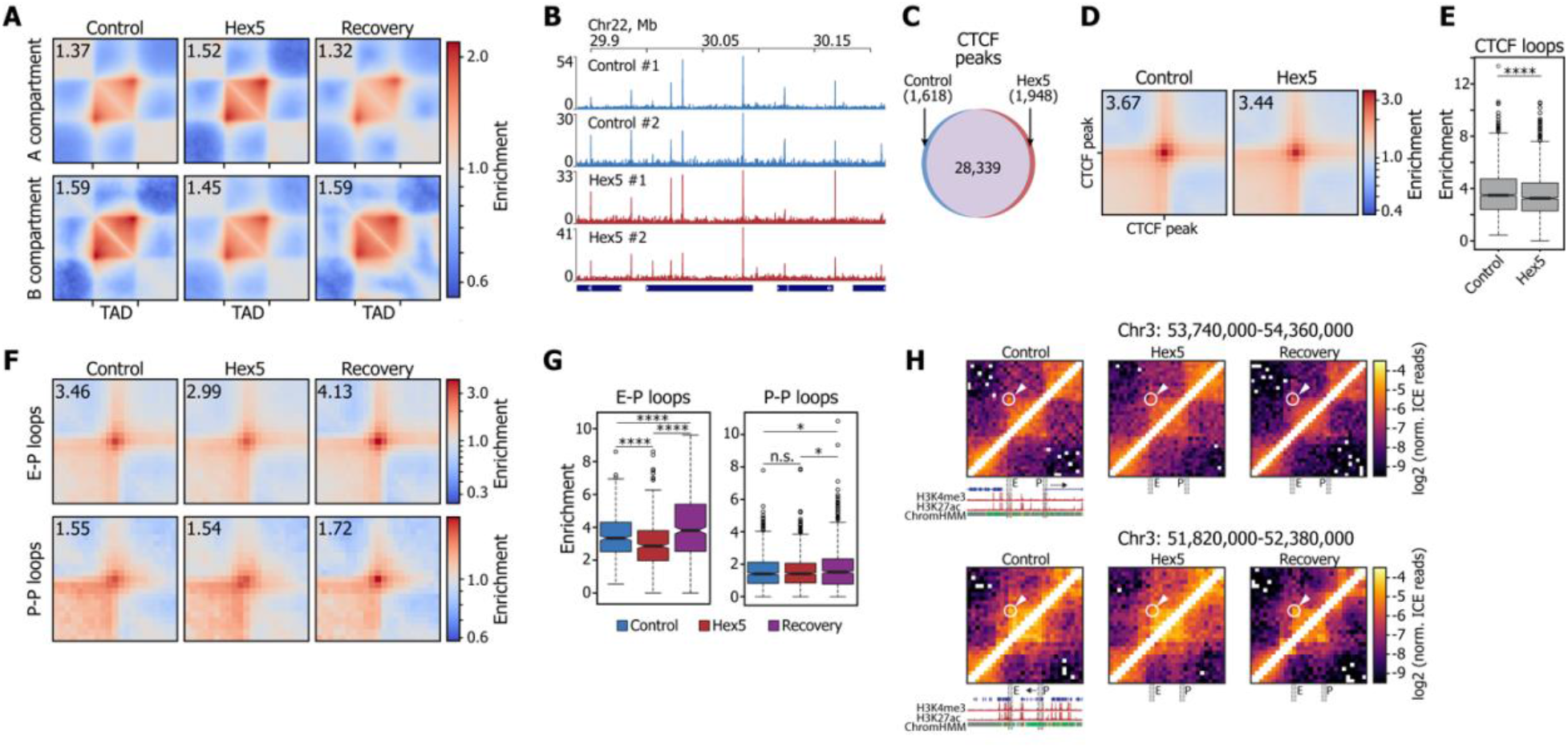
1,6-HD treatment reversibly changes TAD compaction and affects enhancer-promoter interactions. (**A**) Average TADs for the Control, Hex5, and Recovery samples. A number in the upper left corner shows the enrichment of contacts inside the TAD square over the background. (**B**) Examples of CTCF-binding profiles in biological replicates from the Control and Hex5 samples. The dark blue rectangles below the profiles schematically show gene positions. (**C**) Venn diagram showing overlapping CTCF peak sets identified in the Control and Hex5 samples. (**D**) Average CTCF-mediated loop. A number in the upper left corner shows the enrichment of contacts inside the loop pixel over the background. (**E**) Related to (**D**): boxplots showing the distribution of signal in the central pixel of the average CTCF loop plot. \*\*\*\**P* < 0.0001 in a Mann-Whitney U-test. (**F**) Average enhancer-promoter (E-P) and promoter-promoter (P-P) loops. A number in the upper left corner shows the enrichment of contacts inside the loop pixel over the background. (**G**) Related to (**F**): boxplots showing the distribution of signal in the central pixel of the corresponding average loop plot. **** *P* < 0.0001, \**P* < 0.05, n.s., non-significant difference in a Mann-Whitney U-test. (**H**) Representative examples of weakened E-P loops following HEX treatment. Loop pixels are highlighted with a circle and arrow. E, enhancer region, P, promoter region. H3K4me3, H3K27ac, chromatin state (ChromHMM), and gene profiles are shown according to the UCSC genome browser. The resolution of the Hi-C maps is 20 kb.

To study the effects of 1,6-HD treatment on spatial communication among genome regulatory elements, we identified enhancer-promoter (E-P) and promoter-promoter (P-P) loops using chromatin state annotation (Ernst and Kellis 2010). Previously, E-P communications have been reported to occur within phase-separated multi-molecular assemblies (Cho et al. 2018; Sabari et al. 2018). However, whether 1,6-HD-mediated LLPS disruption can affect preformed interactions that were potentially stabilized by other mechanisms, such as specific protein-protein interactions between transcription factors, remains unclear (Deng et al. 2014). Our analysis demonstrated that 1,6-HD treatment significantly reduced the strength of E-P loops (*P* = 6.43×10-8, Mann-Whitney U-test) but not of P-P loops (*P* = 0.36, Mann-Whitney U-test) (Fig. 5E). In the Recovery sample, both types of interactions became stronger compared with those in the Control sample. The reasons for the differential sensitivity to 1,6-HD treatement between the E-P and P-P loops are currently unclear. Interactions among IDR-containing proteins may result in both liquid phase separation and gelation, depending on the conditions (Harmon et al. 2017; St George-Hyslop et al. 2018). Although the liquid condensates formed by LLPS are sensitive to 1,6-HD, the sensitivity of gels to this agent remains unknown. Thus, the data obtained potentially indicate that E-P and P-P interaction have different natures.

## Discussion

Recent evidence suggests that LLPS may contribute to the spatial organization of the eukaryotic genome, particularly to the assembly of the inactive chromatin compartment (Larson et al. 2017; Turner et al. 2018; Tatavosian et al. 2019) and to the mediation of E-P contacts (Hahn 2018). However, most observations related to the role played by LLPS in chromatin folding have either been indirect or based on *in vitro* experiments (Larson et al. 2017; Strom et al. 2017; Boehning et al. 2018; Gibson et al. 2019; Plys et al. 2019; Wang et al. 2019). Here, we used the LLPS compromising agent, 1,6-HD (Sabari et al. 2018), to directly investigate the role played by phase separation in the shaping of the 3D genome in living cells. We found that under conditions that were sufficient to affect Cajal bodies, splicing speckles and HP1α foci, and prevent the formation of optoDroplets, LLPS suppression did not lead to the significant changes in chromatin folding. These results indicate that phase separation sensitive to 1,6-HD is not a key determinant of higher-order genome organization. However, our results suggested that LLPS partially contributes to chromatin compaction at both the local (nucleosome nanodomains) and whole-chromosome (A/B compartments) levels. As revealed by STORM analysis, 1,6-HD-mediated compromising of LLPS in a living cells resulted in the enlargement and more uniform distribution of nanodomains over the nuclear space. This result is in agreement with recently published data showing that short artificial nucleosome arrays can form liquid phase-separated droplets, both under physiological conditions *in vitro*, and when injected into the cell nucleus (Gibson et al. 2019). However, 1,6-HD treatment did not destroy the nanodomains, suggesting the participation of other factors, such as electrostatic interactions (Sun et al. 2005) and macromolecule crowding (Hancock 2007), in the formation of nucleosome assemblies. Thus, our data suggest that, in contrast to some types of protein assemblies (Courchaine et al. 2016; Banani et al. 2017; Boeynaems et al. 2018; Gomes and Shorter 2019), nucleosome aggregates are stabilized by LLPS only to a certain degree.

Megabase-scale chromatin compartmentalization was generally retained upon 1,6-HD treatment. Moderate compartment mixing and only partial decompaction of the B compartment was observed following the transient affection of LLPS, suggesting that the spatial segregation of active and repressed chromatin is a stable feature of the 3D genome organization that persists under various influences such as LLPS disruption, hyperosmotic shock (Amat et al. 2019) or heat shock (Ray et al. 2019).

In contrast to the changes caused by either hyperosmotic (Amat et al. 2019) or hypoosmotic (Golov et al. 2015) shocks, the moderate changes in chromatin folding caused by 1,6-HD-mediated LLPS alteration were not fully reversible. We found that the removal of 1,6-HD from the culture medium did not allow chromatin to adopt its initial configuration, but rather led to further compartmental mixing and the more dense packaging of the B compartment as compared to control cells. E-P interactions also weakened upon 1,6-HD treatment and became stronger than in control cells after 1,6-HD removal. Along with the observations made using high-resolution microscopy, these results suggest that in a living cell, 1,6-HD-sensitive LLPS is one of the 3D genome-shaping forces. The compromising of LLPS is likely to release certain constraints imposed by a variety of biological processes like for example an LLPS-driven assembly of RNA polymerase II-Mediator complexes. These constraints cannot be fully reestablished within the short-term recovery period examined here. Therefore, during the Recovery experiment, chromatin adopted a novel configuration.

In conclusion, our results demonstrated that 1,6-HD-sensitive LLPS is not the primary force supporting local and global spatial genome organization but rather contributes to the fine-tuning of the 3D genome. The possibility that LLPS may be more important for 3D genome assembly under certain conditions (for example, after mitosis) and the relationships between LLPS and the other chromatin-shaping factors deserve further consideration.

## ONLINE METHODS

### Cell culture and treatments

Human HeLa cells were obtained from ATCC. All cells were grown in Dulbecco’s Modified Eagle’s Medium (DMEM), supplemented with 10% fetal calf serum. To obtain transiently permeabilized cells, HeLa cells were incubated with 1% Tween 20 in Dulbecco’s phosphate-buffered saline (DPBS) for 10 min at room temperature, washed twice with DPBS, and transferred to culture medium. To obtain 1,6-HD-treated cells, transiently-permeabilized HeLa cells were incubated with 5% 1,6-HD (Sigma-Aldrich, #240117) in culture medium at room temperature, washed twice with DPBS, and transferred to culture medium. For recovery experiments, HeLa cells that were treated with 1,6-HD were incubated in culture medium for 1.5 h at 37°C. The number of caspase-3/7-positive cells was measured using CellEvent Caspase-3/7 Green Detection Reagent (Invitrogen), according to the manufacturer’s instructions.

### Measurement of transcriptional activity

For 5-ethyniluridine (EU) incorporation, cells were incubated with 200 μM EU (Jena Bioscience) for 15 min at 37°C. After this incubation, cells were washed three times with PBS and fixed in 100% cold (−20°C) methanol for 10 min before staining. The cells were washed three times with PBS and processed using a Click-iT EU Imaging Kit (Life Technologies), according to the manufacturer’s recommendations. The integrated intensities of EU fluorescence were analyzed using CellProfiler software.

### Immunofluorescence

For immunostaining, cells were grown on microscope slides. All samples were fixed in 100% cold methanol (−20°C) for 10 min. After washing in PBS, cells were pre-incubated with 1% bovine serum albumin (BSA) in PBS, with 0.05% Tween 20, for 30 min and were then incubated with antibodies (anti-coilin, Abcam, #ab87913; anti-H2B, Active Motif, #61037; anti-H3K9me3, Active Motif, #39161) in PBS, supplemented with 1% BSA and 0.05% Tween 20, for 1 h at room temperature. After the incubation, cells were washed three times with PBS, supplemented with 0.2% BSA and 0.05% Tween 20. The primary antibodies bound to antigens were visualized using Alexa Fluor 488- or Alexa Fluor 594-conjugated secondary antibodies. The DNA was counterstained with 4,6-diamino-2-phenylindole (DAPI) for 10 min at room temperature. The samples were mounted using Dako fluorescent mounting medium (Life Technologies). The immunostained samples were analyzed using a Zeiss AxioScope A.1 fluorescence microscope (objectives: Zeiss N-Achroplan 40 × /0.65 and EC Plan-Neofluar 100 × /1.3 oil; camera: Zeiss AxioCam MRm; acquisition software: Zeiss AxioVision Rel. 4.8.2; Jena, Germany). The images were processed using ImageJ software (version 1.44).

Samples for Structured Illumination Microscopy (SIM) were mounted in Dako fluorescent mounting medium (Life Technologies) and examined using a Nikon N-SIM microscope (100×/1.49 NA oil immersion objective, 488 nm and 561 nm diode laser excitation). Image stacks (z-steps of 0.2 μm) were acquired with EMCCD camera (iXon 897, An-dor, effective pixel size 60 nm). Exposure conditions were adjusted to get a typical yield about 5000 max counts (16-bit raw image) while keeping bleaching minimal. Image acquisition, SIM image reconstruction and data alignment were performed using NIS-Elements (Nikon)

### OptoDroplet analysis

HeLa cells were transfected using XFect reagent with the pHR-FUSN-mCh-Cry2WT plasmid (Shin et al. 2017), which was a gift from Clifford Brangwynne (Addgene plasmid # 101223). OptoDroplet formation was observed using a Nikon Eclipse Ti-E inverted fluorescence microscope equipped with a Nikon Intensilight C-HGFI light source. Acquisitions were performed using the 100× objective, with the TexasRed filter set for the visualization of the mCherry signal and the FITC filter set for Cry2 activation.

### STORM sample preparation and image acquisition

To perform immunostaining, the culture medium was aspirated, and the cells were washed with 1 × PBS once and fixed in 100% methanol for 10 min at −20°C. After washing once with 1× PBS, the cells were incubated with H2B antibodies diluted in blocking buffer (1% BSA in 1 × PBS with 0.05% Tween 20), at 4°C overnight. The cells were washed 3 times with 1× PBS, for 5 min per wash, and the Alexa Fluor 647-conjugated secondary antibody in the blocking buffer was added to the sample for 1 h, protected from light. The cells were washed three times with 1 × PBS and stored in 1 × PBS before imaging. Immediately before imaging, the buffer was replaced with STORM imaging buffer, containing 10% (w/v) glucose (Sigma-Aldrich), 0.56 mg/ml glucose oxidase (Sigma-Aldrich), 0.17 mg/ml catalase (Sigma-Aldrich), and 0.14 M β-mercaptoethanol (Sigma-Aldrich). All imaging experiments were performed using a commercial STORM microscope system from Nikon Instruments (NSTORM). Laser light at 647 nm was used to excite Alexa Fluor 647. The emitted light was collected by an oil immersion 100×, 1.49 NA objective and imaged onto an electron-multiplying charge-coupled device (EMCCD) camera. For STORM imaging, 40,000 frames were acquired at an exposure time of 20 ms. The reconstruction of the super-resolution image was performed using NanoJ-SRRF Fiji’s plugin.

### STORM image analysis

#### Spatial descriptive statistics

Spatial analysis of H2B was performed as previously described (Nozaki et al. 2017). Briefly, for each extracted nucleus from a reconstructed STORM image, the bottom 10% of signal intensities were discarded as background. Next, we separated each nucleus into 50×50 px squares. For each square, we calculated L- and G-functions, using a custom python code (available at GitHub page: https://github.com/ArtemLuzhin/STORM-microscopy-analysis). As a control, we shuffled the values in each square and then calculated the L- and G-functions for the shuffled squares. The L- and G-function plots were obtained as the ratio between the L-function and the G-function for both raw and shuffled squares.

#### Analysis of nanodomains

The identification and analysis of H2B nanodomains were performed as previously described (Ricci et al. 2015). For each nucleus, a constant threshold was used to digitize the density maps into binary images, such that pixels with a density above the threshold value are given a value of 1, whereas all other pixels are given a value of 0. Binary images were then used to locate regions of the sample that contained localizations, and further analyses were performed on the raw localization data. Localizations falling on zero-valued pixels in the binary images (low-density areas) were discarded from further analyses. To identify H2B nanodomains, we used a K-means clustering algorithm. First, we identify the local maxima of the density map and the cluster-adjacent local maxima. Next, the coordinates for the local maxima cluster centers were used as the initial H2B cluster centers. Localizations were associated with clusters based on their proximity to cluster centroids. New cluster centroid coordinates were iteratively calculated as the average of localization coordinates belonging to the same cluster. The procedure was iterated until the sum of the squared distances between localizations and the associated clusters converged, providing cluster centroid positions and the number of localizations per cluster. Cluster sizes were calculated as the standard deviation (SD) of localization coordinates from the relative cluster centroid.

### Hi-C library preparation

Hi-C libraries were prepared as described previously (Ulianov et al. 2016), with minor modifications. A total of 5-10 million cells were fixed in 1× PBS containing 2% formaldehyde (Sigma-Aldrich) for 10 min, with occasional mixing. The reaction was quenched with by the addition of 2 M glycine (Sigma-Aldrich), to a final concentration of 125 mM. Cells were pelleted by centrifugation (1,000 × g, 10 min, 4 °C), resuspended in 50 μl 1× PBS, snap-frozen in liquid nitrogen, and stored at −80°C. Cells were lysed in 1.5 ml isotonic buffer [50 mM Tris-HCl pH 8.0 (Sigma-Aldrich), 150 mM NaCl (Sigma-Aldrich), 0.5% (v/v) NP-40 substitute (Fluka), 1% (v/v) Triton-X100 (Sigma-Aldrich), 1× Halt™ Protease Inhibitor Cocktail (Thermo Scientific)], on ice for 15 min. Cells were pelleted by centrifugation at 2,500 × g for 5 min, resuspended in 100 μl 1× DpnII buffer (New England Biolabs), and pelleted again. The pellet was resuspended in 200 μl 0.3% SDS (Sigma-Aldrich) in 1.1× DpnII buffer (New England Biolabs) and incubated at 37°C for 1 h. Then, 330 μl 1.1× DpnII buffer and 53 μl 20% Triton X-100 were added, and the suspension was incubated at 37°C for 1 h. Next, 600 U DpnII enzyme (New England Biolabs) were added, and the chromatin was digested overnight (14-16 h), at 37°C, with shaking (1,400 rpm). In the morning, 200 U DpnII enzyme was added, and the cells were incubated for 2 h. DpnII was then inactivated by incubation at 65°C for 20 min. The nuclei were harvested for 10 min at 5,000 × g, washed with 100 μl 1× NEBuffer 2 (New England Biolabs), and resuspended in 125 μl 1.2× NEBuffer 2. Cohesive DNA ends were biotinylated by adding 25 μl biotin fill-in mixture [0.025 mM dATP (Thermo Scientific), 0.025 mM dGTP (Thermo Scientific), 0.025 mM dTTP (Thermo Scientific), 0.025 mM biotin-14-dCTP (Invitrogen), 0.8 U/μl Klenow enzyme (New England Biolabs)]. The samples were incubated at 37°C for 75 min, with shaking (1,400 rpm). Nuclei were pelleted by centrifugation at 3,000 × g for 5 minutes, resuspended in 300 μl 1× T4 DNA ligase buffer (Thermo Scientific), and pelleted again. The pellet was resuspended in 300 μl 1× T4 DNA ligase buffer, and 75 U T4 DNA ligase (Thermo Scientific) was added. Chromatin fragments were ligated at 20°C for 6 h. The cross-links were reversed by overnight incubation at 65°C in the presence of proteinase K (100 μg/ml) (Sigma-Aldrich). After cross-link reversal, the DNA was purified by single phenol-chloroform extraction, followed by ethanol precipitation [glycogen (Thermo Scientific) at a concentration of 20 μg/ml was used as co-precipitator]. After precipitation, the pellets were dissolved in 100 μl 10 mM Tris-HCl, pH 8.0. To remove residual RNA, samples were treated with 50 μg RNase A (Thermo Scientific) for 45 min at 37°C. To remove residual salts and DTT, the DNA was purified using Agencourt AMPure XP beads (Beckman Coulter). Biotinylated nucleotides from the non-ligated DNA ends were removed by incubating the Hi-C libraries (2 μg) in the presence of 6 U T4 DNA polymerase (New England Biolabs) in NEBuffer 2, supplemented with 0.025 mM dATP and 0.025 mM dGTP, at 20°C for 4 h. Next, the DNA was purified using Agencourt AMPure XP beads. The DNA was then dissolved in 500 μl sonication buffer (50 mM Tris-HCl, pH 8.0, 10 mM EDTA, 0.1% SDS) and was sheared to a size of approximately 100-1,000 bp, using a VirSonic 100 (VerTis). The samples were concentrated (and simultaneously purified), using AMICON Ultra Centrifugal Filter Units (Merck), to a total volume of approximately 50 μl. The DNA ends were repaired by adding 62.5 μl MQ water, 14 μl 10× T4 DNA ligase reaction buffer, 3.5 μl 10 mM dNTP mix (Thermo Scientific), 5 μl 3 U/μl T4 DNA polymerase (New England Biolabs), 5 μl 10 U/μl T4 polynucleotide kinase (New England Biolabs), and 1 μl 5 U/μl Klenow DNA polymerase (New England Biolabs) and incubating at 20°C for 30 min. The DNA was purified with Agencourt AMPure XP beads and eluted with 50 μl 10 mM Tris-HCl (pH 8.0). To perform an A-tailing reaction, the DNA samples were supplemented with 6 μl 10× NEBuffer 2, 1.2 μl 10 mM dATP, 1 μl MQ water and 3.6 μl 5 U/μl Klenow (exo-) (New England Biolabs). The reactions were performed for 30 min at 37°C in a PCR machine, and the enzyme was then heat-inactivated by incubation at 65°C for 20 min. The DNA was purified using Agencourt AMPure XP beads and eluted with 200 μl 10 mM Tris-HCl (pH 8.0). Biotin pulldown of the ligation junctions was performed as described previously, with minor modifications. Briefly, 10 μl MyOne Dynabeads Streptavidin C1 (Invitrogen) beads were used to capture the biotinylated DNA, and the volumes of all buffers were decreased by 4-fold. The washed beads attached to captured ligation junctions were resuspended in 50 μl adapter ligation mixture, composed of 41.5 μl MQ water, 5 μl 10× T4 DNA ligase reaction buffer (Thermo Scientific), 2.5 μl Illumina TruSeq adapters and 1 μl 5 U/μl T4 DNA ligase (Thermo Scientific). Adapter ligation was performed at 22°C, for 2.5 h, and the beads were washed twice with 100 μl TWB (5 mM Tris-HCl, pH 8.0, 0.5 mM EDTA, 1 M NaCl, 0.05% Tween-20 (Sigma-Aldrich)), once with 100 μl 1× binding buffer (10 mM Tris-HCl, pH 8.0, 1 mM EDTA, 2 M NaCl), and once with 100 μl CWB (10 mM Tris-HCl, pH 8.0 and 50 mM NaCl), and then resuspended in 20 μl MQ water. Test PCR reactions containing 4 μl streptavidin-bound Hi-C library were performed to determine the optimal number of PCR cycles necessary to generate sufficient PCR products for sequencing. The PCR reactions were performed using KAPA High Fidelity DNA Polymerase (KAPA) and Illumina PE1.0 and PE2.0 PCR primers (10 pmol each). The temperature profile was 5 min at 98°C, followed by 6, 9, 12, 15, and 18 cycles of 20 s at 98°C, 15 s at 65°C, and 20 s at 72°C. The PCR reactions were separated on a 2% agarose gel supplemented with ethidium bromide, and the number of PCR cycles necessary to obtain a sufficient amount of DNA was determined based on the visual inspection of gels (typically 9-12 cycles). Four preparative PCR reactions were performed for each sample. The PCR mixtures were combined, and the products were purified using Agencourt AMPure XP beads.

### Hi-C data processing

Hi-C reads were mapped to the reference human genome hg19 assembly, using Bowtie v2.2.3 (Langmead and Salzberg 2012), with the ‘very-sensitive’ mode and the iterative mapping procedure, implemented in *hiclib* (https://bitbucket.org/mirnylab/hiclib), as described previously (Imakaev et al. 2012). The minimal read size was set to 25 bp, and the iterative mapping step was increasing by 5 bp until a maximal read length was reached. We then filtered out non-uniquely mapped reads, ‘same fragment’ and ‘dangling end’ reads, PCR duplicates, reads from restriction fragments shorter than 100 bp and longer than 100 kb, and reads from the top 0.5% of restriction fragments with the greatest number of reads. The remaining read pairs were aggregated into 20-kb and 500-kb genomic bins to produce high- and low-resolution contact matrices. To remove low-coverage bins and iteratively correct the contact matrices, we used the balance function from *cooler* v0.7.9 (Abdennur and Mirny 2019), with default parameters. Because the two biological replicates in all experiments demonstrated high correlations (Pearson’s *R* > 0.92), they were pooled together for downstream analyses. Statistics for the Hi-C data processing can be found in Supplemental Tables S1 and 3.

### Compartment profile

#### Compartment annotation

Chromatin compartments were annotated using the principal component analysis (PCA) implemented in *mirnylib* (https://bitbucket.org/mirnylab/mirnylib). To identify compartments using 500-kb-resolution contact matrices, we generated the observed-over-expected cis matrix for each chromosome, converted it into a correlation matrix, and partitioned the matrix into eigenvectors by PCA (Lieberman-Aiden et al. 2009). For 20-kb-resolution matrices, we computed eigenvectors directly from the mean-centered observed-over-expected matrix (Imakaev et al. 2012). The eigenvector that represented the compartment signal was selected according to the eigenvalue and correlations with RNA-seq, DNAse-seq, and ChIP-seq results, which were used to validate the eigenvector decomposition. According to the established convention, the orientation of the eigenvector correlated positively with GC content. Consequently, the B-compartment bins were those with negative compartment signals, whereas the A compartment bins were those with positive signals.

#### Validation of the compartment signal

To validate the annotated compartment signals, we used epigenetic feature data for the HeLa cell line from ENCODE (Davis et al. 2018). Raw RNA-seq reads from two biological replicates were mapped to the reference human genome hg19 assembly, using STAR v2.6.1c (Dobin et al. 2013), with default parameters, and then merged. Unmapped and low mapping quality reads were then removed using SAMtools v1.5 (Li et al. 2009), with option -q 30. We calculated the transcription levels in the 20- and 500-kb genomic bins, using BEDtools v2.25.0 (Quinlan and Hall 2010). Aligned DNAse-seq reads from three biological replicates were merged, filtered, and binned, using the same procedure that was applied to RNA-seq data, to obtain chromatin accessibility levels. ChIP-seq signals were aggregated into 20- and 500-kb genomic bins, using BEDtools v2.25.0 (Quinlan and Hall 2010). The statistical significance of epigenetic feature distribution across compartments A and B was determined by a two-sided Mann-Whitney U-test.

### Compartment analysis

#### Saddle plots

Compartmentalization saddle plots were generated as described previously (Imakaev et al. 2012). We used 500-kb observed-over-expected contact matrices, generated by *mirnylib* and *cooltools* (https://github.com/mirnylab/cooltools), for *cis* and *trans* matrices, respectively. In each observed-over-expected matrix, we rearranged the rows and the columns by increasing the compartment signal. We then aggregated the resulting matrices into 25 equally sized bins. These binned matrices were averaged over all chromosomes to obtain cis and trans saddle plots.

#### Compartment strength

Compartment strengths were calculated separately for compartments A and B for each chromosome, as described previously (Falk et al. 2019), using 500-kb-resolution intra- and interchromosomal contact matrices. Compartment strengths were calculated on a per-bin level as the ratio between the average number of contacts the bin makes with bins of the same compartment type and the average number of contacts the bin makes with any bin in the observed-over-expected contact matrix. To obtain per-chromosome compartment strengths we averaged the per-bin strength values for the A and B compartments. Statistical differences between compartment strengths for A and B were determined using the Wilcoxon signed-rank test.

#### Average compartment

To follow the distribution and density of contacts in the A and B compartments, we developed and implemented the average compartment visualization and applied it to the 500-kb-resolution contact matrices. The average compartment visualization, referred to here as a pentad, consisted of five piled-up area types from the observed-over-expected Hi-C matrix, which represented the short- and long-range contacts within the A and B compartments and the contacts between them (Fig. 4A). The areas were determined based on the annotated compartment signals. We considered only the areas that had both dimensions greater than 1 and a maximum number of zero fractions less than 0.1 and were closer than 0.75 times the chromosome length to the diagonal. These areas were extracted from the matrix and rescaled into 33×33-pixel squares, using bilinear interpolation. Rescaled areas of the same type were averaged genome-wide, using the median value for each pixel, and aggregated into one pentad.

### TAD detection

TADs were annotated on 20-kb-resolution matrices, using the Armatus (Filippova et al. 2014) algorithm implementation from the lavaburst package (https://github.com/nvictus/lavaburst). In this algorithm, the average size and the number of TADs are controlled by the scaling parameter, *γ*, which was set to 0.3 to obtain the domain median size of 600 kb (Dixon et al. 2012) for the control experiment. The same *γ* value was then used to identify TADs in other experiments. TADs smaller than 60 kb were excluded due to their poor resolution. 4,828, 3,793, and 6,378 TADs were called in Control, Hex5 and Recovery samples respectively.

### TAD analysis

#### Assignment to the compartments

The annotated TADs were assigned to compartment A or B if the domain intersection with the compartment was greater than 90%. Using this procedure, we assigned 3,024 TADs to compartment A and 583 to compartment B for the control sample, 1,372 to A and 1,031 to B for the 1,6-HD-treated sample, and 4,501 to A and 545 to B for the Recovery sample.

#### Average TAD

The average TADs for compartments A and B were calculated using *coolpup.py* v0.8.6 (Ilya et al. 2019), with the rescale and local options, on 20-kb contact matrices. Domains smaller than 1 Mb were used for pile-up generation (2,933 TADs from compartment A and 327 from compartment B for the control sample, 1,256 from A and 742 from B for the 1,6-HD-treated sample, and 4,425 from A and 338 from B for the Recovery sample) because longer TADs may be artifactual in nature (Dixon et al. 2012; Rao et al. 2014). The rescale size was set to 99 pixels, and 25 randomly shifted control regions were used to normalize the signal for local background. The mean values of the central 33×33-pixel squares that represented the average TAD density (or the enrichment of contacts in the TAD) are highlighted in the top left corner of the pile-ups.

### Loop detection

Loops were annotated using the CPU version of HiCCUPS (Juicer Tools v1.11.09) (Durand et al. 2016), on 5-, 10- and 25-kb-resolution contact matrices, using the recommended parameters for medium resolution Hi-C maps. Annotations from different resolution loops were merged, as described previously (Rao et al. 2014). 2,837, 2,381, and 5,709 loops were annotated Control, Hex5 and Recovery samples, respectively.

### Loop analysis

#### CTCF-associated loops

To identify CTCF-associated loops, we used the annotated CTCF ChIP-seq peaks and HiCCUPS loops for control and 1,6-HD-treated samples. A loop with at least one base intersecting with a ChIP-seq peak was annotated as being CTCF-associated. CTCF loops with separations of 0.1-1.5 Mb were used for the average loop construction (2,563 for control and 2,065 for 1,6-HD-treated cells).

#### Enhancer-promoter interactions

To identify E-P loops we utilized the ENCODE combined chromatin state segmentation for the HeLa cell line. We searched for loops that had a ‘TSS’ (transcription start site) state in one loop base and either an ‘E’ (enhancer) or ‘WE’ (weak enhancer) state in the other, as described previously (Rao et al. 2014). The identified E-P loops with separations of 0.1-1.5 Mb were used for the average loop construction (455 loops).

#### Promoter-promoter interactions

Gene pairs with P-P interactions were detected by ChIA-PET, and promoter coordinates were detected by CapStarr-seq for the HeLa cell line, which were obtained from a recent publication (Dao et al. 2017). Interacting promoters with separations of 0.1-1.5 Mb were used for the average loop construction (929 pairs).

#### Average loop

The average loop was calculated using *coolpup.py* v0.8.6 (Ilya et al. 2019), on 20-kb contact matrices, with a pad size of ± 200 kb around the loop pixel, and 10 randomly shifted control regions to normalize the signal for local background. The value in the central pixel shows the average loop strength (or the enrichment of contacts in the loop) and is highlighted in the top left corner of the pile-up.

### Interchromosomal contacts

The proximity of the chromosome territories was calculated as the ratio between the observed and expected number of interactions between chromosomes *i* and *j*, as described previously (Lieberman-Aiden et al. 2009). The expected number of contacts between chromosomes *i* and *j* was computed by multiplying the fraction of total interchromosomal contacts containing chromosome *i* with the fraction of total interchromosomal contacts containing chromosome *j* and multiplying by the total number of interchromosomal contacts.

### *P_c_*(*s*) curves

*P_c_*(*s*) curves were computed using *hiclib*, and the range between 20 kb and 100 Mb was taken.

### Downsampling of contacts

We performed downsampling of the obtained Hi-C matrices using the previously described approach (Yardimci et al. 2019). The Hi-C matrix was converted into a set of pairwise interactions from which we uniformly sampled a given number of contacts. These downsampled contacts were then re-binned into the Hi-C matrix at a 500 kb resolution.

### ChIP-seq library preparation

ChIP-seq was performed with an anti-CTCF antibody (Active Motif, #61311) as described (Pena-Hernandez et al. 2015; Arrigoni et al. 2016) for two biological replicates. ChIP samples were prepared for next-generation sequencing using a NEBNext Ultra II DNA library prep kit for Illumina (New England Biolabs). Libraries were sequenced on the Illumina NovaSeq 6000 and resulted in around 40 million 100-bp single-end reads per sample.

### CTCF ChIP-seq data analysis

Reads were mapped to the reference human genome hg19 assembly, using Bowtie v2.2.3 (Langmead and Salzberg 2012), with the ‘very-sensitive’ mode. Non-uniquely mapped reads and possible PCR and optical duplicates were filtered out using SAMtools v1.5 (Li et al. 2009). The bigWig files, with the ratio of RPKM normalized ChIP-seq signal to the input, were generated using *deepTools2* (Ramirez et al. 2016) for each biological replicate. Peaks were called using PePr (Zhang et al. 2014), with a p-value cutoff of 0.05 and a sliding window size of 100 bp. The peak calling procedure annotated 29,957 peaks in control and 30,287 peaks in 1,6-HD-treated sample. The annotated peaks were assigned to compartment A or B if the peak completely intersected with the compartment. The heatmaps of the ChIP-seq signal, centered at the peak coordinates, were generated using *deeptools2* (Ramirez et al. 2016).

### Data availability

The accession numbers of the publicly available data used in this work are indicated in Supplemental Table S2. Raw sequencing reads for the Hi-C and ChIP-seq libraries and the processed Hi-C matrices, annotated compartments, TADs and loops, ChIP-seq signals and peaks are available in the GEO repository, under accession number GSE138543.

## Acknowledgements

This work was supported by the Russian Science Foundation (19-74-10009). We thank the Center for Precision Genome Editing and Genetic Technologies for Biomedicine, IGB RAS for providing the equipment to perform Hi-C data analysis. SIM and STORM experiments were performed using the equipment purchased within the framework of Lomonosov Moscow State University development program (PNR 5.13).

## Author Contributions

O.L.K. and S.V.R. conceived the study; S.V.U. performed Hi-C experiments; A.K.V. performed cell culturing, immunostaining and transcription analysis; M.M.D, A.V.L. and A.V.T. analyzed publicly available, Hi-C and ChIP-seq data; N.O. and I.I.K. performed SIM and STORM; A.V.L. perfomed SIM and STORM data analysis; A.K.G. and A.A.G prepared ChIP-seq libraries; O.L.K., S.V.U. and S.V.R. wrote the manuscript with input from all co-authors.

## SUPPLEMENTAL FIGURE LEGENDS

**Supplemental Figure S1. 1,6-HD affects LLPS in living human cells.** (**A**) Quantification of the samples presented in Fig. 1D. Percentage of cells containing coilin foci (i.e. Cajal bodies) are shown.

(**B**) HeLa cells were first transiently permeabilized with Tween 20 and then were either left untreated (control) or treated with 1,6-HD (5%, 15 min) before being stained for SC35 (SRSF2). The DNA was stained with DAPI. The samples were analyzed by structured illumination microscopy (SIM). Nanodomain clustering analysis (see Methods for details) was performed (13 control cells and 9 1,6-HD-treated cells were analyzed). Average nanodomain cluster sizes and numbers of nanodomains per cell are depicted in each case. (**C**) HeLa cells treated as in (**B**) were stained for histone H2B and analyzed by epifluorescence microscopy. The DNA was stained with DAPI. Scale bar: 10 μm.

**Supplemental Figure S2. Visualization of Control, Hex5, and Recovery contact matrices at a 20-kb resolution.**

**Supplemental Figure S3. Hi-C data processing and compartment annotation.** (**A**) Pearson’s correlation coefficient between biological replicates of the Hi-C experiments. (**B**) Heat map of the interchromosomal contacts in Control, Hex5, and Recovery samples. Each bin of the map corresponds to one chromosome (labeled). (**C**) Boxplots showing the levels of different epigenetic markers in the A and B compartments, annotated in control cells. (**D**) Scatter plots showing the correlations among PC1 values for Control, Hex5, and Recovery samples. (**E**) Coverage of the genome with A and B compartments. (**F**) Distributions of PC1 values in Control, Hex5, and Recovery samples.

**Supplemental Figure S4. Reproducibility of Hi-C data chrematistics between biological replicates.** Dependence of contact probability, *P_c_*(*s*), on genomic distance, *s* (**A**) and ratio between *cis* (intrachromosomal) and *trans* (interchromosomal) contacts (**B**) for Control, Hex5, and Recovery biological replicates.

**Supplemental Figure S5. Hi-C data processing and compartment annotation in control experiments (Tween-treated HeLa cells).** (**A**) Pearson’s correlation coefficient between biological replicates of the Hi-C experiments. (**B**) Ratio between *cis* (intrachromosomal) and *trans* (interchromosomal) contacts. (**C**) Visualization of Control, Tween, and Recovery contact matrices at a 500-kb resolution. (**D**) Coverage of the genome with A and B compartments. (**E**) Distributions of PC1 values in Control, Tween, and Recovery samples.

**Supplemental Figure S6. Reanalysis of the Hi-C data reported by Amat et al.** (**A**) Examples of Hi-C matrices visualized at a 500-kb resolution. (**B**) Compartment strength in control cells, after hyperosmotic shock (NaCl), and after recovery. (**C**) Plots (pentads) showing the observed over the expected contact frequencies inside the A and B compartments at short and large genomic distances and between compartments. (**D**) Pairwise subtractions of Control, NaCl, and Recovery pentads. (**E**) Boxplots showing the contact frequencies between the A and B compartments.

**Supplemental Figure S7. CTCF ChIP-seq analysis.** (**A**) Pearson’s correlation coefficient between biological replicates of the ChIP-seq experiments. (**B**) Heat map of the CTCF ChIP-seq signal at CTCF peaks, identified using the PePr tool.

## SUPPLEMENTAL TABLE LEGENDS

**Supplemental Table S1.** Statistics of the Hi-C data processing.

**Supplemental Table S2.** List of publicly available data used in this work.

**Supplemental Table S3.** Statistics of the Hi-C data processing in control experiments (Tween-treated HeLa cells).

